# Obesity-associated lipidomic remodeling of the adrenal gland indicates an important role of the FADS2-arachidonic acid axis in adrenocortical hormone production

**DOI:** 10.1101/2020.09.04.282905

**Authors:** Anke Witt, Peter Mirtschink, Alessandra Palladini, Ivona Mateska, Heba Abdelmegeed, Michal Grzybek, Ben Wielockx, Mirko Peitzsch, Ünal Coskun, Triantafyllos Chavakis, Vasileia Ismini Alexaki

## Abstract

**Objective:** Adrenocortical hormone levels increase in obesity, potentially contributing to development of obesity-associated pathologies. Here we explored whether lipidomic remodeling of the adrenal gland could mediate altered adrenocortical steroidogenesis during obesity.

**Methods:** Lipidomic analysis was performed in adrenal glands using shotgun mass spectrometry (MS), and steroid profiling of sera by liquid chromatography tandem mass spectrometry (LC-MS/MS) from lean and obese mice. Gene expression analysis was performed in adrenal glands and adrenocortical cell populations. The role of Fatty Acid Desaturase 2 (FADS2) and arachidonic acid on steroid hormone production was studied in primary adrenal gland cell cultures.

**Results:** Adrenal glands of obese mice displayed a distinct lipidomic profile, encompassing longer and more unsaturated storage lipids and phospholipids compared to adrenal glands of lean mice. Arachidonoyl acyl chains were abundant in the adrenal gland phospholipidome and increased upon obesity. This was accompanied by increased *Fads2* expression, the rate-limiting enzyme of arachidonic acid synthesis, and enhanced plasma adrenocortical hormone levels. Inhibition of FADS2 in primary adrenal gland cell cultures abolished steroidogenesis, which was restored by arachidonic acid supplementation.

**Conclusions:** Our data suggest that the FADS2 – arachidonic acid axis regulates adrenocortical hormone synthesis, while alterations in the content of arachidonoyl chains in the adrenal gland phopsholipidome could account for disturbed adrenocortical hormone production.

**Highlights:**

- The adrenal gland lipidome is remodeled in obesity.
- Arachidonoyl groups are abundant in the adrenal gland phospholipidome and increase in obesity.
- FADS2 is highly expressed in the adrenal gland and its expression is further increased in obesity.
- FADS2 inhibition blunts adrenocortical steroidogenesis in primary adrenal gland cell cultures, while arachidonic acid supplementation restores it.

## 1. Introduction

Obesity increases the risk for type 2 diabetes mellitus, non-alcoholic steatohepatitis and cardiovascular diseases [1, 2]. Elevated adrenal hormone levels are observed in obesity and could contribute to the development of obesity-associated pathologies [3–8]. Aldosterone, the major regulator of blood volume and pressure [9], is increased in the circulation in obesity, which can cause hypertension and cardiovascular disease [4, 6, 7, 10, 11]. Circulating glucocorticoids, which can promote abdominal fat deposition, are also elevated in animal models of obesity [4, 6, 12]. Moreover, elevated aldosterone and glucocorticoid levels can promote insulin resistance [7, 13, 14]. However, the mechanisms underlying adrenal function alterations during obesity remain largely obscure.

A feature of obesity is dysregulation of lipid homeostasis in tissues, such as the adipose tissue and liver, resulting in accumulation of storage lipids, i.e. triacylglycerides (TAG) in these tissues [15–18]. Moreover, the chain length and saturation of lipid species in the adipose tissue and liver, as well as the serum lipidome substantially change during obesity [16, 17, 19–22]. In the present study we investigated how the lipid landscape of the adrenal gland changes upon diet-induced obesity and whether these changes are involved in the regulation of adrenal hormone synthesis. We identified arachidonic acid as an important phospholipid component in the adrenal gland. The rate-limiting enzyme of arachidonic acid synthesis, FADS2, is highly expressed in the adrenal gland and its expression is up-regulated in the adrenal glands of obese animals. Moreover, FADS2 inhibition in primary adrenal gland cell cultures abolished adrenocortical hormone production and responsiveness to ACTH stimulation, while arachidonic acid supplementation restored it. These data suggest an important role of the FADS2-arachidonic acid axis in adrenal hormone steroidogenesis.

## 2. Materials and Methods

### 2.1 Feeding experiments

Six-weeks-old male C57BL/6 mice were fed a normal diet (ND, D12450B, Research Diets, NJ, USA), or a high fat diet (HFD, D12492, Research Diets, NJ, USA) for 20 weeks, as previously described [23–25]. At the end of the feeding the mice were sacrificed and organs were isolated. Experiments were approved by the Landesdirektion Sachsen, Germany.

### 2.2 Lipid extraction for mass spectrometry lipidomics

Adrenal glands were homogenized in ammonium-bicarbonate buffer (150 mM ammonium bicarbonate, pH 7) with TissueLyser (Qiagen). Protein content was assessed using BCA Protein Assay Kit (Thermo Fisher). Equivalents of 20 μg of protein were taken for mass spectrometry (MS) analysis. MS-based lipid analysis was performed by Lipotype GmbH (Dresden, Germany) as described [17, 26]. Lipids were extracted using a two-step chloroform (Sigma Aldrich) / methanol (Thermofisher Scientific, Darmstadt, Germany) procedure [27]. Samples were spiked with internal lipid standard mixture containing: cardiolipin 16:1/15:0/15:0/15:0 (CL), ceramide 18:1;2/17:0 (Cer), hexosylceramide 18:1;2/12:0 (HexCer), lyso-phosphatidate 17:0 (LPA), lyso-phosphatidylcholine 12:0 (LPC), lyso-phosphatidylethanolamine 17:1 (LPE), lyso-phosphatidylglycerol 17:1 (LPG), lyso-phosphatidylinositol 17:1 (LPI), lyso-phosphatidylserine 17:1 (LPS), phosphatidate 17:0/17:0 (PA), phosphatidylcholine 17:0/17:0 (PC), phosphatidylethanolamine 17:0/17:0 (PE), phosphatidylglycerol 17:0/17:0 (PG), phosphatidylinositol 16:0/16:0 (PI), phosphatidylserine 17:0/17:0 (PS), cholesterol ester 20:0 (CE), sphingomyelin 18:1;2/12:0;0 (SM), cholesterol D6 (Chol) (all Avanti Polar Lipids), triacylglycerol 17:0/17:0/17:0 (TAG) and diacylglycerol 17:0/17:0 (DAG) (both Larodan Fine Chemicals). Synthetic lipid standards were purchased from Avanti Polar Lipids, Larodan Fine Chemicals and Sigma–Aldrich and all chemicals were analytical grade. After extraction, the organic phase was transferred to an infusion plate and dried in a speed vacuum concentrator (Martin Christ, Germany). The dry extract from the first step was re-suspended in 7.5 mM ammonium acetate (Sigma Aldrich) in chloroform/methanol/propan-2-ol (Thermofisher Scientific) (1:2:4 V:V:V) and the second-step dry extract in 33 % ethanol of methylamine (Sigma Aldrich)/ chloroform/methanol (0.003:5:1 V:V:V) solution. All liquid handling steps were performed using Hamilton Robotics STARlet robotic platform with the Anti Droplet Control feature for organic solvents pipetting.

### 2.3 MS data acquisition

Samples were analyzed by direct infusion on a QExactive mass spectrometer (Thermo Fisher Scientific) equipped with a TriVersa NanoMate ion source (Advion Biosciences, Ithaca, NY). Samples were analyzed in both positive and negative ion modes with a resolution of R_(m/z=200)_=280000 for MS and R_(m/z=200)_=17500 for MS/MS experiments, in a single acquisition. MS/MS was triggered by an inclusion list encompassing corresponding MS mass ranges scanned in 1 Da increments [28]. Both MS and MS/MS data were combined to monitor CE, DAG and TAG ions as ammonium adducts; PC, PC O-, as acetate adducts; and CL, PA, PE, PE O-, PG, PI and PS as deprotonated anions. MS only was used to monitor LPA, LPE, LPE O-, LPI and LPS as deprotonated anions; Cer, HexCer, SM, LPC and LPC O- as acetate adduct and cholesterol as ammonium adduct of an acetylated derivative [29].

### 2.4 Data processing and downstream analysis

Data were analyzed with an in-house developed lipid identification software based on LipidXplorer [30, 31]. Data post-processing and normalization were performed using an in-house developed data management system. Only lipid identifications with a signal-to-noise ratio > 5 and a signal intensity 5-fold higher than in corresponding blank samples were considered for further data analysis. All downstream analyses were performed in R (R Core Team 2019). The dataset was filtered by applying an occupational threshold of 50%, meaning that lipids that were present in less than 50% of the samples in a group were set to zero in that group. Dataset was log2-scaled before Principal Component Analysis (PCA). The descriptive analysis was performed on the mol %-transformed dataset, i.e. picomol quantities were divided by the sum of the lipids detected in the respective sample and multiplied by 100. Total carbon chain length and double bonds plots result from grouping together all the lipids that present the same number of carbon atoms (total length) or the same number of double bonds (degree of unsaturation) and calculating the mean and standard deviation in each group of samples (ND and HFD). To further inspect general trends in the degree of unsaturation and length of storage and membrane lipids under different perturbations, the average weighted number of double bonds (Double Bond Index, DBI) and length (Length Index, LI), as previously described [32]. Statistical tests were performed on the mol %-transformed data. First, normality was assessed by means of the Shapiro-Wilkinson test, and according to its result either the Welch t-test or the Wilcoxon test was used. P-values were then adjusted following the Benjamini-Hochberg correction. Significance was set at p<0.05. Analysis was done in R [33]. Plots were generated using ggplot2 [34].

### 2.5 Steroid hormones measurements

Steroid hormones in plasma and cell culture supernatants were analyzed by liquid chromatography tandem mass spectrometry (LC-MS/MS) as described elsewhere [35]. In brief, 50 to 100μL of mice plasma or cell culture supernatants were extracted by solid phase extraction using positive pressure, followed by a dry down under an gentle stream of nitrogen. Residues were reconstituted in 100 μL of initial LC mobile phase and 10 μL were injected into the LC-MS/MS. Quantification of steroid concentrations was done by comparisons of ratios of analyte peak areas to respective peak areas of stable isotope labeled internal standards obtained in samples to those of calibrators.

### 2.6 Sorting of adrenocortical cell populations

Adrenal glands were isolated and cleaned from the surrounding fat tissue and the adrenal cortex was separated from the medulla under a dissecting microscope. The cortex was collected in a digestion buffer, consisting of collagenase I and bovine serum albumin (both at 1.6 mg/ml concentration, Sigma-Aldrich, Germany) dissolved in phosphate-buffered saline (PBS), and digested for 25 minutes at 37 °C while shaking at 900 rpm. Dissociated cells were passed through a 22 G needle and then a 100 μm cell strainer and centrifuged at 300 x g for 5 minutes at 4 °C. Cells were then washed in MACS buffer (0.5 % BSA, 2 mM EDTA in PBS) and endothelial cells (CD31^+^) and leukocytes (CD45^+^) were sequentially positively selected using CD31 and CD45 MicroBeads (Miltenyi Biotec), respectively, according to manufacturer’s instructions. Briefly, pelleted cells were resuspended in 190 μl MACS buffer, mixed with 10 μl CD31 MicroBeads, incubated for 15 minutes at 4 °C, washed with 2 ml MACS buffer and centrifuged at 300 x g for 10 minutes. The cell pellet was resuspended in 500 μl MACS buffer and applied onto MS Column placed on MACS Separator. The columns were washed and the flow-through was collected. CD31^+^ cells were positively sorted. The flow-through was centrifuged at 300 x g for 5 minutes, and the pelleted cells were subjected to the same procedure using CD45 MicroBeads. The flow-through containing CD31^−^CD45^−^ adrenocortical cells was centrifuged at 300 x g for 5 minutes, and the pelleted cells were collected. CD45^+^ cells were positively sorted. Collected cell populations were kept for transcriptional analysis.

### 2.7 Primary adrenal cell culture and treatment

Adrenal glands were isolated from wild-type C57BL/6 mice, they were cleaned from the surrounding fat tissue and incubated in collagenase buffer (1.6 mg/ml collagenase type I, 1.6 mg/ml BSA) for 45 minutes at 37 °C, while shaking at 900 rpm. Dissociated cells were passed through 22 G needle and a 100 μm cell strainer and centrifuged at 300 x g for 5 minutes at 4 °C. Cells of a pool of two adrenals originating from one mouse were distributed in four 0.2 % gelatin-precoated wells of a 96-well plate in DMEM/F12 supplemented with 1 % FBS, 50 U/mL penicillin and 50 μg/mL streptomycin. Two hours after seeding 10 μM FADS2 inhibitor (Tocris, Wiesbaden-Nordenstadt, Germany) or same amount of carrier (dimethylsulfoxide, DMSO) were added without changing the medium. In some experiments medium was additionally supplemented with 150 μM arachidonic acid (Sigma Aldrich, Germany) or same amount of solvent (endotoxin-free water). In some experiments, the medium was changed 18 hours later and the cells were incubated with or without 10 ng/mL Adrenocorticotropic hormone (ACTH) in the presence of either 10 μM Fads2 inhibitor or DMSO for 60 minutes. Supernatants were collected for steroid hormone measurement and cell lysates were collected for transcriptional analysis.

### 2.8 Transcriptional analysis

RNA was isolated from frozen tissues with the TRI Reagent (MRC, Cincinnati, USA) after mechanical tissue disruption or from cells with the RNeasy Mini Kit (Qiagen, Düsseldorf, Germany) according to manufacturer’s instructions. RNA obtained with Tri Reagent was subsequently extracted with Chloroform and the NucleoSpin RNA, Maxi kit (Macherey-Nagel, Düren, Germany). Reverse transcription was performed with the iScript cDNA Synthesis kit (Biorad, Munich, Germany) and cDNA was analyzed by qPCR using the SsoFast Eva Green Supermix (Bio-Rad, Munich, Germany), a CFX384 real-time System C1000 Thermal Cycler (Bio-Rad) and the Bio-Rad CFX Manager 3.1 software. Primers used are listed in Supplementary Table 1.

### 2.9 Statistical analysis

Data were analyzed with a Mann-Whitney U test or a one-way ANOVA with post-hoc Tukey’s test for multiple comparisons with p<0.05 set as a significance level using the GraphPrism 7 software.

## 3. Results

### 3.1 Adrenal glands from lean and obese mice presented distinct lipidomic profiles

We first asked whether obesity affects the adrenal gland lipidome. To this end, mice were fed for 20 weeks a ND or HFD and shotgun lipidomics was performed in isolated adrenal glands. HFD-feeding led to a significant increase of the body weight (not shown) [24, 25, 36]. PCA was performed on the whole lipidome (Figure 1A), non-storage lipids, i.e. phospholipids and their ether-linked versions (PC, PE, PC O-, PE O-, PI, PG, PS, PA), lysolipids (LPA, LPC, LPE, LPE O-, LPG, LPI, LPS), sphingolipids (SM), DAG and cardiolipins (Figure 1B), and storage lipids (CE and TAG) (Figure 1C) [17, 37–40]. Whole and non-storage lipidomes of adrenal glands from lean and obese mice segregated along the first principal component (PC1) and the topology of the samples was strikingly similar (Figure 1A,B), while storage lipids regrouped with statistical significance along the second principal (PC2) (Figure 1C), suggesting that non-storage lipids were mainly responsible for the grouping of the whole lipidome according to the diet.

**Figure 1.**
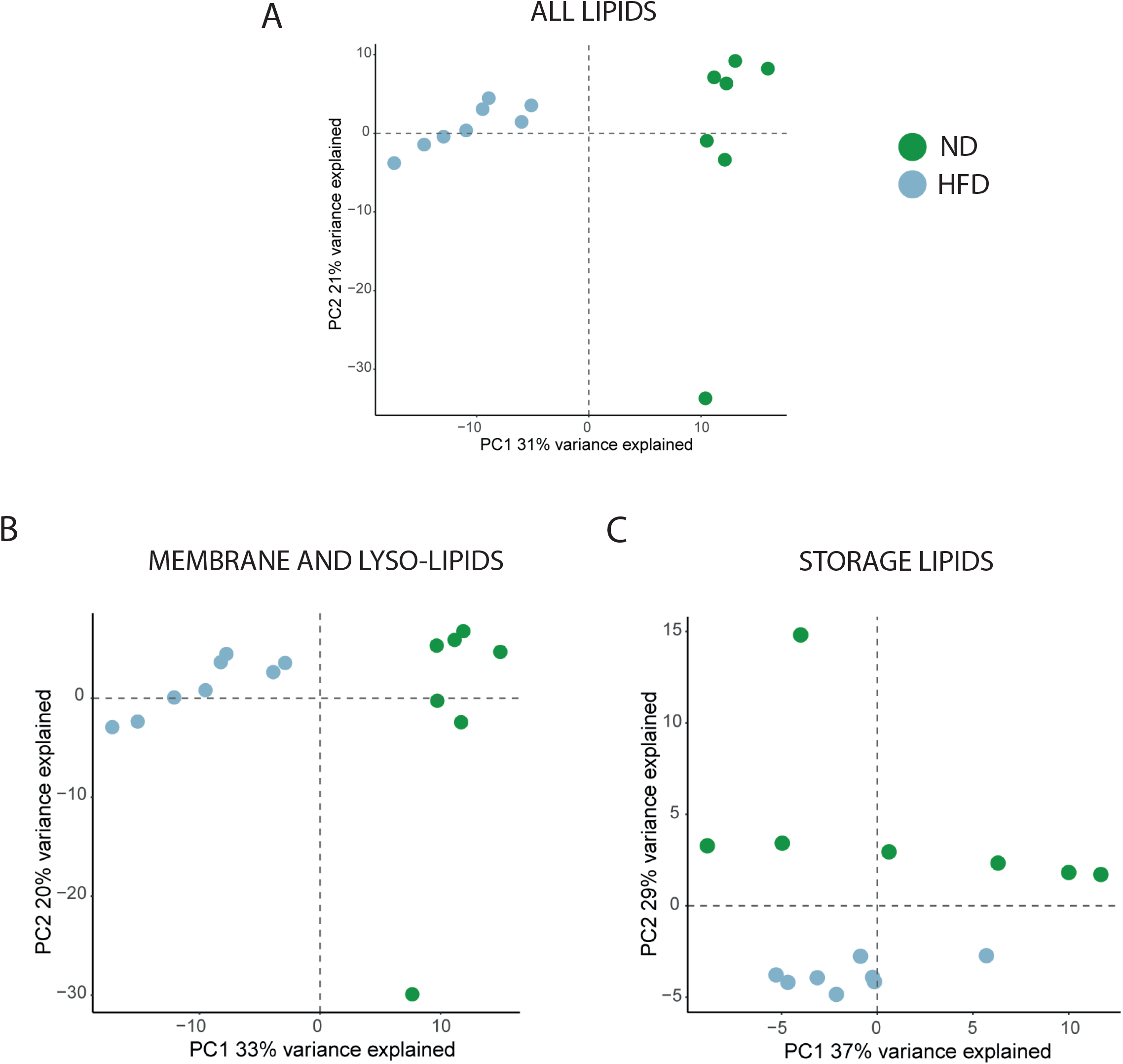
Principal component analysis on lipids of adrenal glands of lean and obese mice. Principal component analysis (PCA) performed on all lipids (A), membrane and lyso-lipids (B) and storage lipids (C) in adrenal glands from mice fed for 20 weeks a ND or a HFD, n=7-8.

Yet, it was storage lipids (TAG and CE) that constituted the largest portion of the identified lipids in the adrenal glands (86.98 ± 5.43 % in mice on ND and 90.12 ± 1.93 % in mice on HFD), while membrane lipids accounted for 12.64 ± 5.22 % and 9.63 ± 1.89 % and lysolipids for 0.38 ± 0.23 % and 0.24 ± 0.06 % of all identified lipids in the adrenal glands of mice under ND and HFD, respectively (Table 1). Hence, the total amounts of storage, membrane lipids and lysolipids did not substantially differ in the adrenal glands of lean and obese mice. Also at the lipid class level there were no significant differences between the adrenal glands of lean and obese mice (Table 2, Supplementary Figure 1). The most abundant non-storage lipid class was PC followed by cholesterol and PE (Table 2, Supplementary Figure 1).

**Table 1.** Lipidome composition of adrenal glands in ND- and HFD-fed mice.

|  | **Storage lipids**<br>**(% mean ± Sd)** | **Membrane lipids**<br>**(% mean ± Sd)** | **Lysolipids**<br>**(% mean ± Sd)** |
| --- | --- | --- | --- |
| **ND** | 86.98 ± 5.43 | 12.64 ± 5.22 | 0.38 ± 0.23 |
| **HFD** | 90.12 ± 1.93 | 9.63 ± 1.89 | 0.24 ± 0.06 |

**Table 2.** Lipid class abundance in adrenal glands from ND- and HFD-fed mice.

| **pmol/μg protein**<br>**(Aver ± SD)** | **ND** | **HFD** |
| --- | --- | --- |
| **CE** | 246.59 ± 68.57 | 280.49 ± 41.86 |
| **Cer** | 0.86 ± 0.16 | 1.27 ± 0.35 |
| **Chol** | 37.03 ± 5.65 | 37.12 ± 3.64 |
| **CL** | 1.35 ± 0.27 | 1.44 ± 0.31 |
| **DAG** | 12.84 ± 8.06 | 10.26 ± 3.26 |
| **HexCer** | 0.0781 ± 0.0186 | 0.0646 ± 0.0089 |
| **LPA** | 0.00430 ± 0.00398 | 0.00628 ± 0.00148 |
| **LPC** | 3.98 ± 1.25 | 3.49 ± 0.47 |
| **LPE** | 0.91 ± 0.2 | 0.84 ± 0.11 |
| **LPE-O** | 0.33 ± 0.12 | 0.28 ± 0.05 |
| **LPG** | 0.0171 ± 0.0053 | 0.0246 ± 0.0075 |
| **LPI** | 0.0876 ± 0.0123 | 0.1330 ± 0.0188 |
| **LPS** | 0.0287 ± 0.0139 | 0.0395 ± 0.0052 |
| **PA** | 0.0198 ± 0.0200 | 0.0304 ± 0.0149 |
| **PC** | 68.54 ± 6.82 | 70.56 ± 12.32 |
| **PC-O** | 2.07 ± 0.15 | 1.95 ± 0.27 |
| **PE** | 24.72 ± 5.09 | 29.84 ± 7.06 |
| **PE-O** | 18.23 ± 2.49 | 17.84 ± 1.65 |
| **PG** | 0.869 ± 0.067 | 0.766 ± 0.406 |
| **PI** | 12.15 ± 1.94 | 13.3 ± 2.76 |
| **PS** | 5.9 ± 0.49 | 5.73 ± 0.72 |
| **SM** | 5.29 ± 0.75 | 5.56 ± 0.75 |
| **TAG** | 1306.32 ± 767.16 | 1583.34 ± 338.58 |
| **Total** | 3496.45 ± 1473.05 | 4128.73 ± 698.88 |

### 3.2 The chain length and saturation of adrenal gland lipids changed upon obesity

Next, the total acyl chain length and degree of unsaturation (number of double bonds) of the adrenal gland storage (CE, TAG) and non-storage lipids was analyzed. TAG species with 50 to 54 carbon atoms (Supplementary Figure 2A) and one to four double bonds (Supplementary Figure 2B) in their acyl chains were highly abundant (mol % > 10). Adrenal glands of HFD-fed mice presented a significant increase in long (with a total acyl chain length of 54 carbon atoms) and unsaturated (with four, five and more than six double bonds) TAG species. In contrast, adrenal glands from ND-fed mice showed a larger amount of TAG species containing shorter acyl chains (with 48 and 50 carbon atoms) with one or two double bonds (Supplementary Figure 2A,B). To identify whether differences between diets affected the overall mean weighted length and unsaturation level of storage and non-storage lipids, the DBI and the LI were calculated. Storage lipids in the adrenal glands of obese mice showed a significantly higher DBI compared to lean mice (Supplementary Table 2). Hence, upon HFD feeding TAG in the adrenal glands became overall longer and more unsaturated.

Similar analysis was conducted for non-storage lipids. Most non-storage lipids contained a cumulative of 34 to 38 carbon atoms (Supplementary Figure 2C) and one, two, four or five double blonds in their acyl chains (Supplementary Figure 2D). Similarly to what was found for TAG, adrenal glands from HFD-fed mice showed a significant increase in longer non-storage lipids compared to adrenal glands from ND-fed mice, while shorter non-storage lipid species were more abundant in the adrenal glands of ND-fed mice (Supplementary Figure 2C). Also in non-storage lipids, the increase in length correlated with an increase in the number of double bonds: lipid species with 4 double bonds were more abundant in the adrenal glands from obese mice, while lipid species with one double bond were more abundant in the adrenal glands from lean mice (Supplementary Figure 2D). In accordance, membrane lipids in the adrenal glands of HFD-fed mice had significantly higher DBI and LI compared to ND-fed mice (Supplementary Table 2). In summary, membrane lipids, similarly to storage lipids, were significantly longer and more unsaturated in the adrenal glands of obese mice.

### 3.3 20:4 groups were abundant in the phospholipidome of the adrenal gland and increased upon obesity

In order to further understand the obesity-associated changes in the lipidome of the adrenal gland, we analyzed the acyl chain composition of non-storage lipids and cholesterol esters. In this analysis, the amount (as mol %) of the acyl groups was calculated within each lipid class. Strikingly, 20:4, was one of the most abundant acyl chain within CE and many phospholipid classes, i.e. PC, PC O-, PE, PE O-, PG and PI (Figure 2). Moreover, the amount of CE, PC, PE and PG lipids containing 20:4 was increased in the adrenal glands of obese compared to lean mice (Figure 2). Other abundant acyl chains were palmitic acid (16:0) in DAG and PC, stearic acid (18:0) in PC, PE, PI and PS, and 18:1 fatty acids in CE, DAG, PG and PS. In contrast, polyunsaturated fatty acids (PUFAs) other than arachidonic acid, such as 22:4, 22:5 and 22:6, were much less abundant than 20:4 in the adrenal gland phospholipidome (Figure 2).

**Figure 2.**
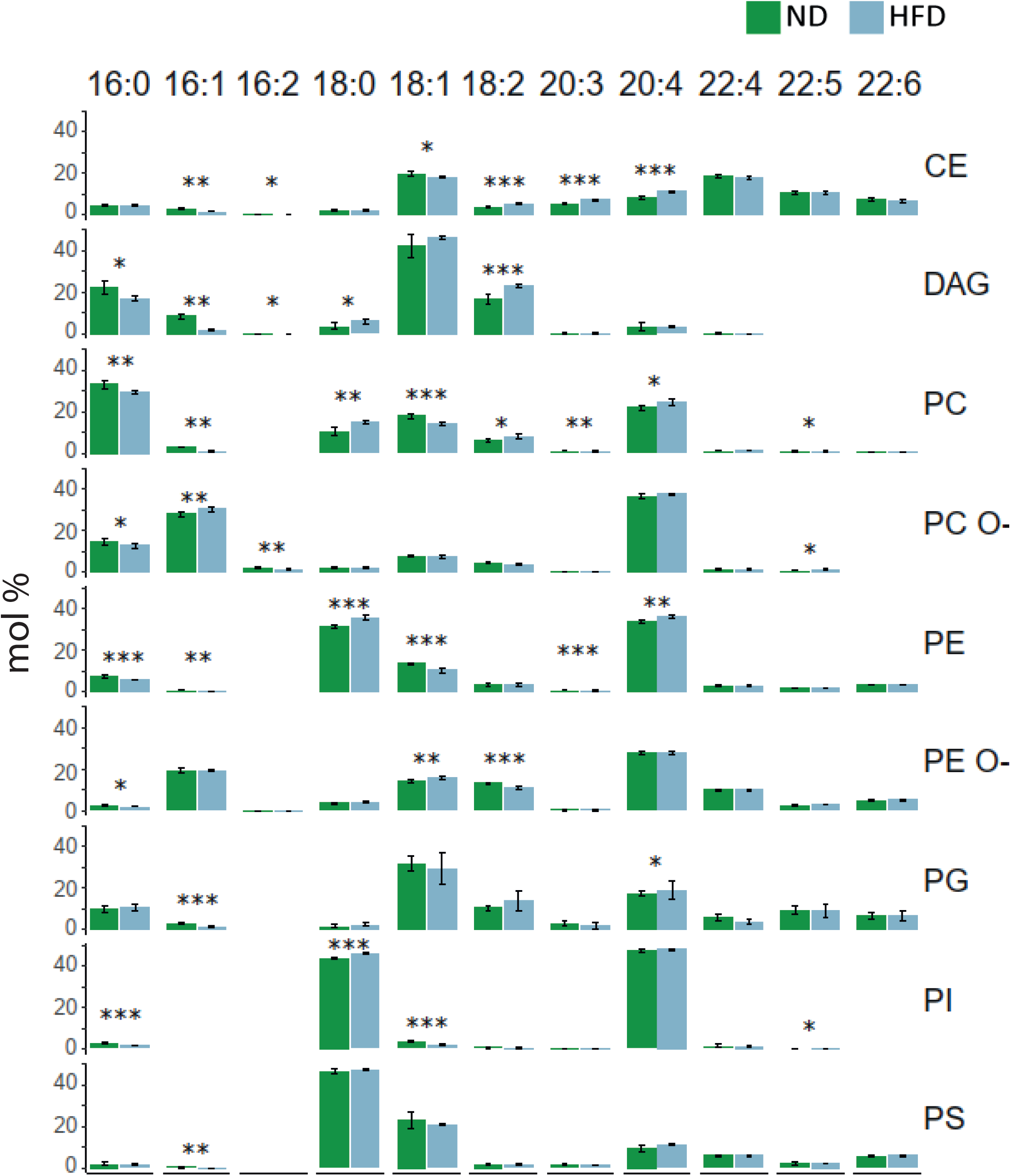
Acyl chain profile of non-storage lipids and cholesterol esters. Lipids in adrenal glands from lean and obese mice were grouped according to the acyl chain within each lipid class (mol % is relative to the lipid class). Only the features with a mean abundance > 5 mol % are shown. The mean mol % ± standard deviation are shown, n=7-8; * p-value<0.05; ** p-value<0.01; *** p-value<0.001.

Next, the specific acyl chain composition of PC and PE, which were both found to be rich in 20:4 (Figure 2), was analyzed. Strikingly, the most abundant PC species were 16:0_20:4 and 18:0_20:4, and the latter increased upon HFD (Figure 3A). In PE, the most abundant species were 18:0_20:4 and these also increased upon HFD (Figure 3B). We hypothesized, that the abundance and increase upon HFD of 20:4 in PC and PE, could indicate a role of arachidonic acid (20:4(n-6), *all-cis-*5,8,11,14-eicosatetraenoic acid) in the adrenal gland function in the lean and obese state. We focused on arachidonic acid and not its isomer 20:4 (n-3) eicosatetraenoic acid (*all*-cis-8,11,14,17-eicosatetraenoic acid), since the latter is, in contrast to arachidonic acid, very little incorporated into the tissue lipidome [41].

**Figure 3.**
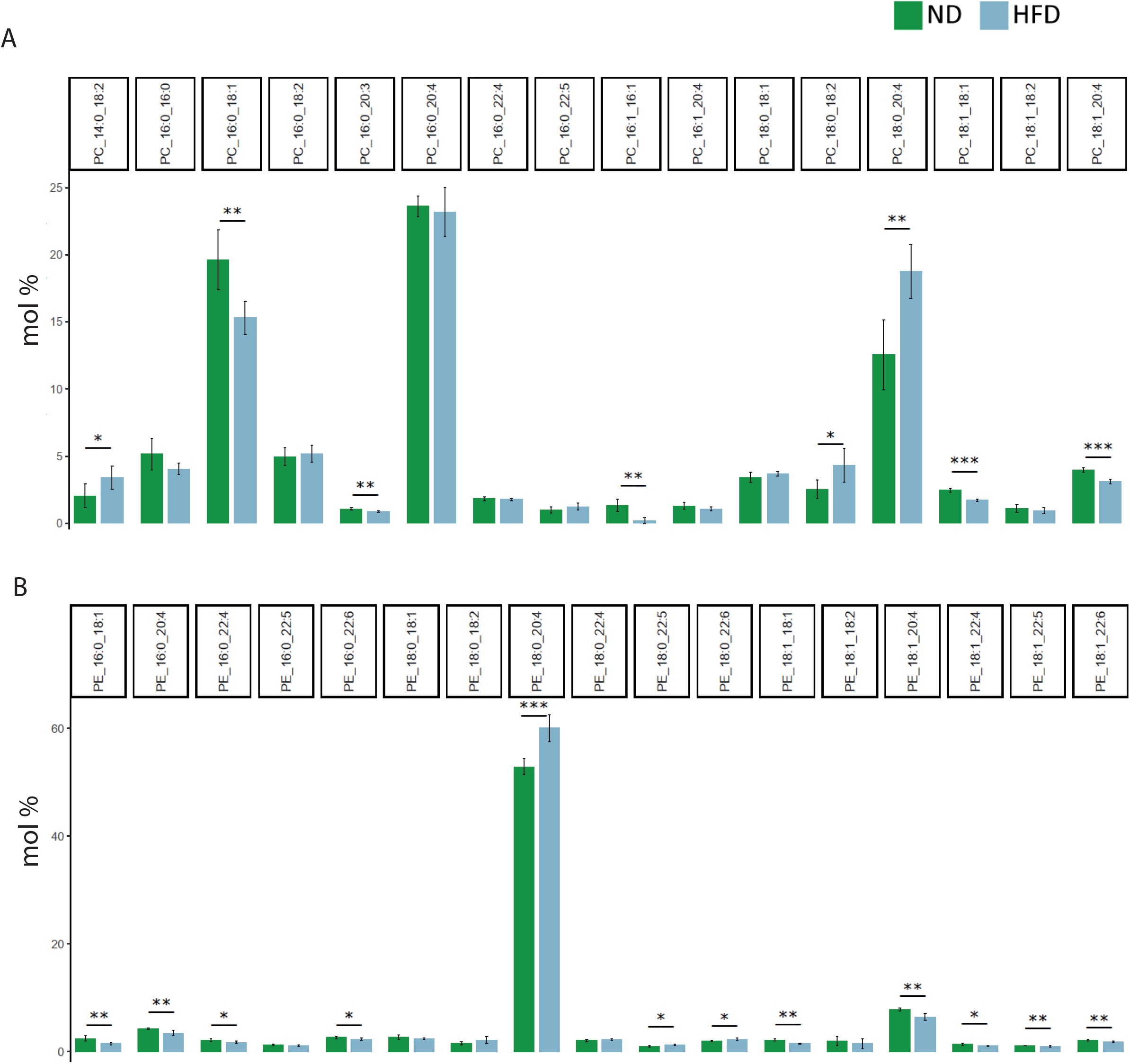
Species composition of PC and PE classes. PC (**A**) and PE (**B**) lipid species in adrenal glands of lean and obese mice. Only the species with a mean abundance > 1 mol % are shown. The mean mol % ± standard deviation are shown, n=7-8; * p-value<0.05; ** p-value<0.01; *** p-value<0.001.

### 3.4 FADS2 was strongly expressed in the adrenal glands and its expression was upregulated upon obesity

We then sought to investigate the mechanism underlying the abundance of 20:4 in the adrenal gland lipidome. Arachidonic acid is endogenously synthesized from the essential fatty acid linoleic acid (*all-cis-*9-12-18:2) and is then esterified to form phospholipids [42–45]. The Δ-6-desaturase FADS2 catalyzes the rate-limiting conversion of linoleic acid to γ-linolenic acid (*all-cis-*6,9,12–18:3) [42–44]. We first examined *Fads2* mRNA expression in the adrenal glands and compared it to different tissues. Following the liver, the adrenal gland was the tissue with the highest *Fads2* expression, followed by the kidney and the brain. Other tissues, such as the spleen, white and brown adipose tissue, lung, heart, skeletal muscle, and testis showed lower *Fads2* mRNA levels (Figure 4A). We also examined *Fads2* expression in different cell populations of the adrenal cortex, i.e. adrenocortical (CD31^−^CD45^−^), endothelial (CD31^+^) cells and leukocytes (CD45^+^) and found that the CD31^−^CD45^−^ adrenocortical cell population displayed the highest *Fads2* expression among these cell populations (Figure 4B).

**Figure 4.**
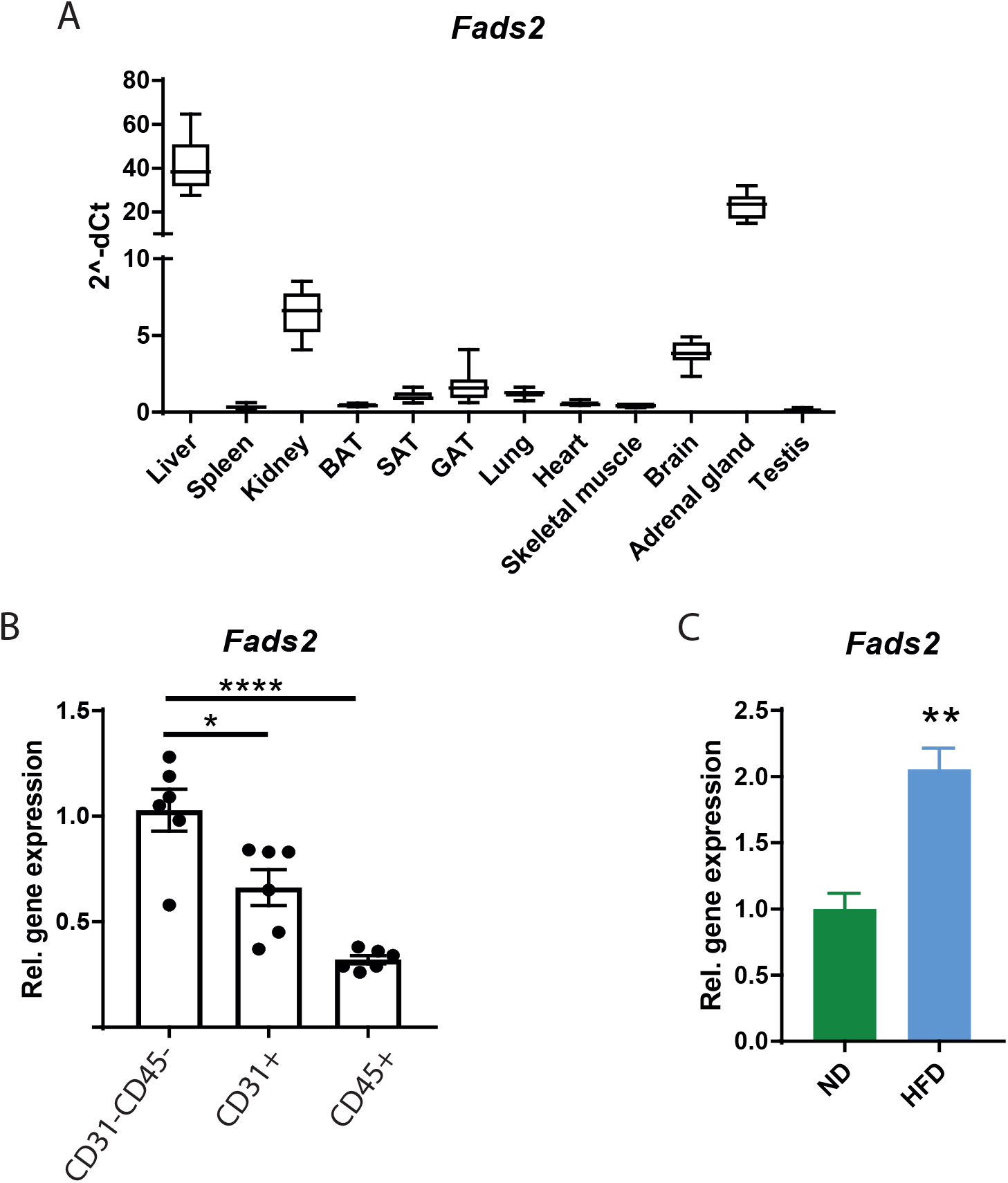
*Fads2* expression in different tissues and adrenocortical cell populations, and the adrenal glands of lean and obese mice. **A.** Relative mRNA level of *Fads2* in different tissues of 8-week old wild-type mice. Data are shown as mean 2^-ΔCt. **B.** Relative *Fads2* expression in CD31^−^CD45^−^, CD31^+^ and CD45^+^ cell populations of the murine adrenal cortex. Data are shown as mean ± SEM, n=6, * p-value<0.05; **** p-value<0.0001. **C.** Relative *Fads2* expression in adrenal glands from lean and obese mice. Expression of *Fads2* in adrenal glands of ND-fed mice was set to 1. Data are shown as mean ± SEM, n=5-6, ** p-value<0.01. For all assays *Tbp<* expression was used as an internal control.

Moreover, *Fads2* expression was increased in the adrenal glands of HFD – fed mice (Figure 4C), which stood in accordance with increased levels of arachidonoyl chains in phospholipids in the adrenal glands of obese mice (Figure 2 and 3). We also asked whether increased *Fads2* expression in obese animals is a unique feature of the adrenal gland. While *Fads2* expression was increased in the liver it was reduced in the white adipose tissue of obese mice (data not shown) suggesting a tissue-specific regulation of *Fads2* expression during obesity. Furthermore, enhanced *Fads2* expression in the adrenal gland of obese animals was associated with elevated plasma levels of corticosterone (ND: 39.08 ± 5.133, HFD: 67.17 ± 10.27; p=0.044) and its precursors, 11-deoxycorticosterone (ND: 0.3236 ± 0.08384, HFD: 1.181 ± 0.3481; p=0.0383) and progesterone (ND: 0.1288 ± 0.02318, HFD: 0.8091 ± 0.2164; p=0.0016).

### 3.5 Adrenocortical hormone production was blocked by FADS2 inhibition and restored by arachidonic acid supplementation

Having identified high *Fads2* expression and abundance of arachidonoyl chains in the adrenal gland, as well as increase of both during obesity correlating with increased adrenocortical steroidogenesis, we set out to investigate whether the FADS2-arachidonic acid axis contributes to adrenal steroidogenesis. To this end, primary adrenal cell cultures were treated with a FADS2 inhibitor and subsequently stimulated with ACTH. ACTH efficiently increased progesterone, 11-deoxycorticosterone, corticosterone and aldosterone production, while FADS2 inhibition abolished the production of all the aforementioned steroid hormones in cells stimulated or not with ACTH (Figure 5A-D). FADS2 inhibition did not affect cell viability (data not shown). Moreover, the effect of FADS2 inhibition on steroidogenesis was not due to decreased mRNA expression of steroidogenic enzymes (*Steroidogenic acute regulatory protein* (*StAR*), *Cytochrome P450 Family 11 Subfamily A Member 1* (*Cyp11a1*, *Cholesterol side-chain cleavage enzyme*), *3b-hydroxysteroid dehydrogenase type 2* (*3b-HSD2*), *Cyp21a1* (*21-hydroxylase*), *Cyp11b1* and *Cyp11b2*). In fact, mRNA expression of all the aforementioned steroidogenic enzymes was increased upon FADS2 inhibition (Supplementary Figure 3), perhaps due to a compensatory response to low steroid hormone synthesis in the cells treated with the FADS2 inhibitor. Finally, to confirm if the inhibition of adrenal steroidogenesis results specifically from decrease in arachidonic acid levels, we examined whether arachidonic acid supplementation could restore adrenocortical hormone production in cultured cells under FADS2 blockage. Indeed, arachidonic acid supplementation counteracted the effect of FADS2 inhibition restoring progesterone, 11-deoxycorticosterone, corticosterone and aldosterone production (Figure 5E-H), suggesting that FADS2-dependent arachidonic acid biosynthesis is required for steroidogenesis in the adrenal gland.

**Figure 5.**
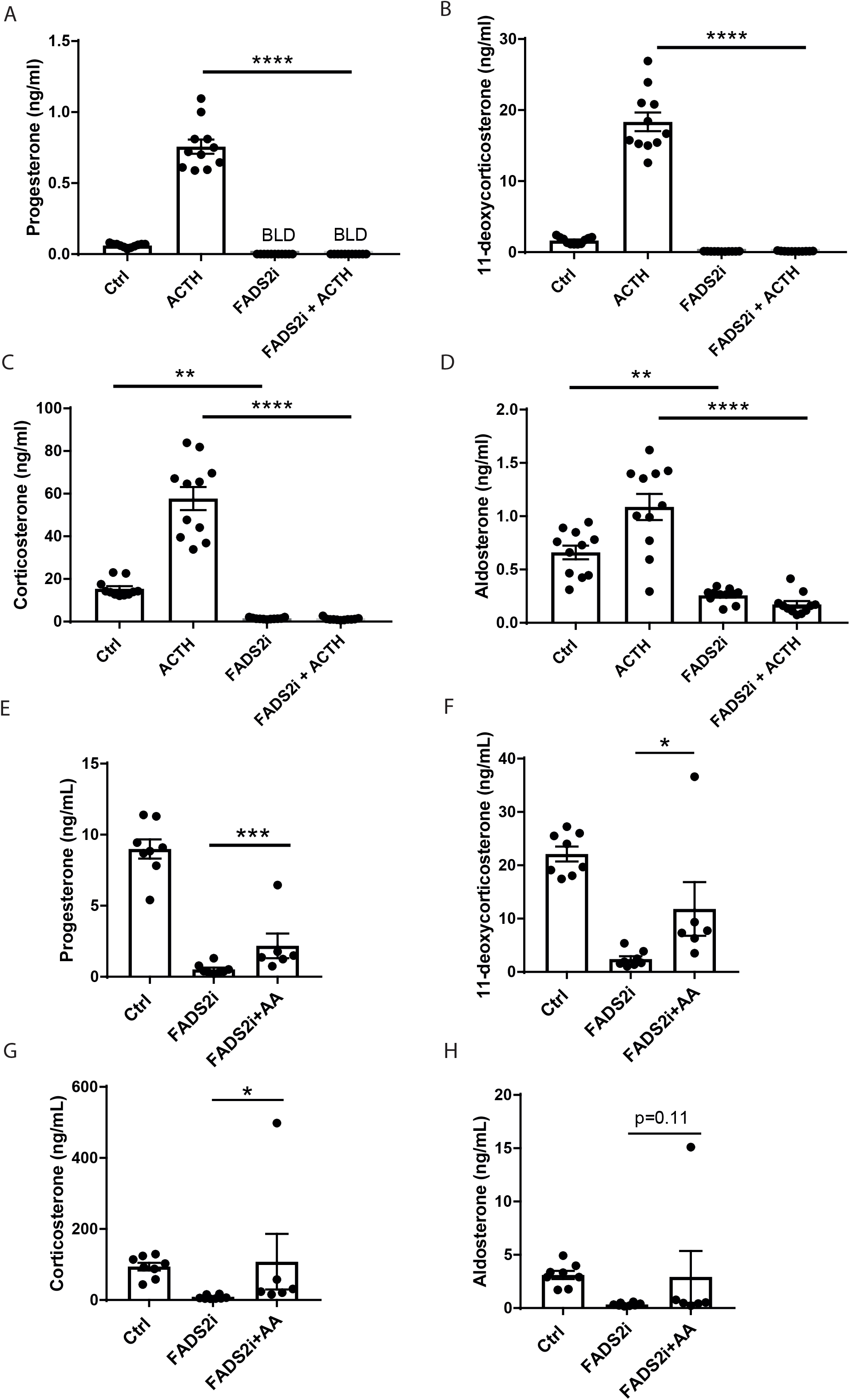
Effect of FADS2 inhibition on adrenocortical hormone production. **A-D.** Progesterone, 11-deoxycorticosterone, corticosterone and aldosterone levels in supernatants of primary adrenal gland cell cultures treated for 18 h with FADS2 inhibitor (FADS2i, 10 μM) or DMSO and then stimulated for 1 h with ACTH (10 ng/mL) or carrier (PBS) in the presence of FADS2 inhibitor or DMSO. The cell medium was changed after the 18 h incubation with FADS2 inhibitor or DMSO. Data are shown as mean ± SEM, np=12, * p-value<0.05; **** p-value<0.0001. BLD – below detection limit. **E-H.** Progesterone, 11-deoxycorticosterone, corticosterone and aldosterone levels in supernatants of primary adrenal gland cell cultures treated for 18 h with FADS2 inhibitor (10 μM) or DMSO in the presence of arachidonic acid (AA) (150 μM) or carrier (endotoxin-free water).

## 4. Discussion

During obesity adrenocortical hormone serum levels rise [3, 4, 6, 46, 47], which contributes to complications such as insulin resistance and hypertension [7, 10, 11, 13, 48]. The molecular mechanisms mediating the elevated adrenocortical hormone production during obesity have been little investigated. Here, we explored such mechanisms by analyzing the lipidomic landscape of the adrenal gland in lean and obese mice, which led to the identification of FADS2 and arachidonic acid as determinants of adrenocortical steroidogenesis.

We found that HFD feeding reshaped the adrenal gland lipidome. Storage and non-storage lipids became longer and more unsaturated upon HFD-feeding, as previously observed in other tissues, such as the adipose tissue and skeletal muscle [17, 49]. Strikingly, 20:4 was among the most abundant acyl chains in CE and several phospholipid classes. This stands in accordance with previous findings, showing high amounts of arachidonic cholesterylesters in rat adrenal glands [50]. 18:0_20:4 PC and PE species were particularly abundant in the adrenal gland lipidome and their levels increased even more upon HFD feeding. PC and PE, which constitute the majority of the eukaryotic phospholipidome, are among the most abundant phospholipids in the endoplasmatic reticulum and mitochondrial membranes [38, 51–54] and are both important stores of arachidonoyl residues [45, 49, 54, 55]. Abundance of PUFA, such as 20:4, in phospholipids constructing phospholipid layers increases membrane fluidity, in contrast to saturated fatty acids, which make membranes more rigid [56, 57]. Hence, membrane fluidity of the adrenal gland cells, encompassing more unsaturated phospholipids and in specific more lipid species with 20:4 acyl chains, might be greater in HFD-fed animals. Membrane fluidity can determine enzymatic activity and secretory functions [39, 56–58]. Steroid hormones are rapidly synthesized from cholesterol through a cascade of enzymatic reactions taking place in the mitochondria and the endoplasmatic reticulum [59]. Moreover, the activity of the enzymes involved in steroid biosynthesis depends on the membrane fluidity of these organelles [60, 61]. Hence, increased membrane fluidity due to higher abundance of arachidonoyl residues in PC and PE could contribute to enhanced adrenocortical hormone synthesis in obese animals. Arachidonic acid predominantly derives from endogenous synthesis from the essential fatty acid linoleate [42, 43, 45]. The first and rate-limiting step of arachidonic acid biosynthesis is the conversion of linoleate to γ-linoleate, which is catalyzed by FADS2 [62, 63]. FADS2 deficiency leads to loss of arachidonoyl chains in the phospholipidome of different tissues including the adrenal glands [62]. Moreover, single nucleotide polymorphisms in the *FADS2* promoter were associated with lower arachidonic levels in serum phospholipids in humans [64]. FADS2 deficiency was reported to lead to a non-canonical conversion of linoleate to eicosatrienoic acid (ω6-20:3), which substituted arachidonic acid in phospholipids thereby critically changing the structure and function of endoplasmatic reticulum and Golgi membranes [65]. Moreover, FADS2 deficient female and male mice are sterile, which is attributed to changes in the membrane fluidity of gonadal cells due to the loss of PUFA acyl groups, which are replaced by shorter and more saturated acyl chains [62, 63], while sterility can be overcome by diet supplementation with arachidonic, eicosapentaenoic and docosahexaenoic acids [62, 66].

We found that FADS2 was highly expressed in the adrenal gland compared to other tissues, such as the lung, heart, brain, spleen, kidney, testis, skeletal muscle and adipose tissue, standing in accordance with a previous report [67]. Moreover, FADS2 expression was regulated during obesity in a tissue-specific manner: in the adrenal glands and liver it increased, while in the adipose tissue it decreased. Along the same line, increased Δ-6-desaturase activity was previously associated with obesity, insulin resistance, steatohepatitis and non-alcoholic fatty liver disease [68–71], while its inhibition protected against liver steatosis and inflammation [72–74].

Arachidonic acid can be converted to the endocannabinoid anandamide, which was previously found to be increased in the adrenal glands of HFD-fed mice [42, 75]. However, gene expression of *Fatty acid amide hydrolase* (*Faah*), the enzyme converting arachidonic acid to anandamide [76, 77], was not altered upon HFD feeding (data not shown). Moreover, the content of lipids in arachidonoyl chains can also be influenced by the release of arachidonic acid from phospholipids via Phospholipase A2 [42, 45]. Among several genes encoding for different Phospholipase A2 isoforms, we found *Pla2g5* and *Pla2g4a* highly expressed in the adrenal gland but their expression was not altered during obesity (data not shown). These findings collectively suggest that the increased content of arachidonoyl chains in the adrenal gland lipidome primarily relies on FADS2-mediated arachidonic acid synthesis.

Finally, we showed that FADS2 inhibition abolished adrenocortical steroidogenesis, which was restored by administration of exogenous arachidonic acid, thereby revealing a decisive role of the FADS2 – arachidonic acid axis in enabling steroidogenesis. This knowledge could be therapeutically harnessed for the management of aberrant adrenocortical hormone production in diseases, such as Cushing’s syndrome or hyperaldosteronism [78, 79].

In conclusion, we demonstrated metabolic disease-driven lipidomic remodeling and particularly alterations in the FADS2-arachidonic acid axis in the adrenal gland. These findings contribute to a deeper understanding of the molecular mechanisms governing adrenocortical hormone production.

## Abbreviations

ACTH: adrenocorticotropic hormone
CE: cholesterol ester
Cer: ceramide
Chol: cholesterol
CL: cardiolipin
DAG: diacylglycerol
DBI: double bond index
FADS2: Fatty Acid Desaturase 2
HexCer: hexosylceramide
HFD: high-fat diet
LC-MS/MS: liquid chromatography tandem mass spectrometry
LI: length index
LPA: lyso-phosphatidate
LPC: lyso-phosphatidylcholine
LPE: lyso-phosphatidylethanolamine
LPE O-: ether-linked LPE
LPG: lyso-phosphatidylglycerol
LPI: lyso-phosphatidylinositol
LPS: lyso-phosphatidylserine
ND: normal diet
PA: phosphatidic acid
PC: phosphatidylcholine
PC O-: ether-linked PC
PCA: principal component analysis
PE: phosphatidylethanolamine
PE O_: ether-linked PE
PG: phosphatidylglycerol
PI: phosphatidylinositol
PS: phosphatidylserine
PUFA: polyunsaturated fatty acid
SM: sphingomyelin
TAG: triacylglyceride
*StAR*: *Steroidogenic acute regulatory protein*
*Cyp11a1*: *Cytochrome P450 Family 11 Subfamily A Member 1*
*3b-HSD2*: *3b-hydroxysteroid dehydrogenase type 2*
*Cyp21a1*: *Cytochrome P450 Family 21 Subfamily A Member 1*
*Cyp11b1*: *Cytochrome P450 Family 11 Subfamily B Member 1*
*Cyp11b2*: *Cytochrome P450 Family 11 Subfamily B Member 2*

## Acknowledgments

We gratefully acknowledge Denise Kaden and Christine Mund for their technical assistance, and Sider Penkov for reviewing the manuscript.

This work was supported by funds from the Deutsche Forschungsgemeinschaft (SFB-TRR 205 to VIA), the ERC (DEMETINL to TC) and the German Federal Ministry of Education and Research grant to the German Center for Diabetes Research (ÜC, TC).

## Declarations of interest

None

## Author contributions

**Anke Witt:** Methodology, validation, formal analysis, investigation, data curation, writing - review & editing, visualization; **Peter Mirtschink:** Conceptualization, methodology, investigation, writing - review & editing; **Alessandra Palladini:** Software, validation, formal analysis, data curation, writing - review & editing; **Ivona Mateska:** Methodology, investigation, writing - review & editing; **Heba Abdelmegeed:** Investigation. **Michal Grzybek:** Data curation, writing - review & editing; **Ben Wielockx:** writing - review & editing; **Mirko Peitzsch:** Methodology, investigation, writing - review & editing; **Ünal Coskun:** Resources, writing - review & editing, funding acquisition; **Triantafyllos Chavakis:** Conceptualization, resources, writing - review & editing, funding acquisition; **Vasileia Ismini Alexaki:** Conceptualization, validation, formal analysis, resources, data curation, writing - original draft, writing - review & editing, visualization, supervision, project administration, funding.

## Notes

### Competing Interest Statement

The authors have declared no competing interest.

## Literature

1. Bluher M. Obesity: global epidemiology and pathogenesis. Nat Rev Endocrinol. 2019;15(5):288–98.

2. Heymsfield SB, Wadden TA. Mechanisms, Pathophysiology, and Management of Obesity. N Engl J Med. 2017;376(3):254–66.

3. Kargi AY, Iacobellis G. Adipose tissue and adrenal glands: novel pathophysiological mechanisms and clinical applications. Int J Endocrinol. 2014;2014:614074.

4. Swierczynska MM, Mateska I, Peitzsch M, Bornstein SR, Chavakis T, Eisenhofer G, et al. Changes in morphology and function of adrenal cortex in mice fed a high-fat diet. Int J Obes (Lond). 2015;39(2):321–30.

5. Reimann M, Qin N, Gruber M, Bornstein SR, Kirschbaum C, Ziemssen T, et al. Adrenal medullary dysfunction as a feature of obesity. Int J Obes (Lond). 2017;41(5):714–21.

6. Hofmann A, Peitzsch M, Brunssen C, Mittag J, Jannasch A, Frenzel A, et al. Elevated Steroid Hormone Production in the db/db Mouse Model of Obesity and Type 2 Diabetes. Horm Metab Res. 2017;49(1):43–9.

7. Roberge C, Carpentier AC, Langlois MF, Baillargeon JP, Ardilouze JL, Maheux P, et al. Adrenocortical dysregulation as a major player in insulin resistance and onset of obesity. Am J Physiol Endocrinol Metab. 2007;293(6):E1465–78.

8. An Y, Reimann M, Masjkur J, Langton K, Peitzsch M, Deutschbein T, et al. Adrenomedullary function, obesity and permissive influences of catecholamines on body mass in patients with chromaffin cell tumours. Int J Obes (Lond). 2019;43(2):263–75.

9. Feldman RD. Aldosterone and blood pressure regulation: recent milestones on the long and winding road from electrocortin to KCNJ5, GPER, and beyond. Hypertension. 2014;63(1):19–21.

10. Calhoun DA, Sharma K. The role of aldosteronism in causing obesity-related cardiovascular risk. Cardiol Clin. 2010;28(3):517–27.

11. Northcott CA, Fink GD, Garver H, Haywood JR, Laimon-Thomson EL, McClain JL, et al. The development of hypertension and hyperaldosteronism in a rodent model of life-long obesity. Endocrinology. 2012;153(4):1764–73.

12. Galitzky J, Bouloumie A. Human visceral-fat-specific glucocorticoid tuning of adipogenesis. Cell Metab. 2013;18(1):3–5.

13. Geer EB, Islam J, Buettner C. Mechanisms of glucocorticoid-induced insulin resistance: focus on adipose tissue function and lipid metabolism. Endocrinol Metab Clin North Am. 2014;43(1):75–102.

14. Hofmann A, Brunssen C, Peitzsch M, Martin M, Mittag J, Jannasch A, et al. Aldosterone Synthase Inhibition Improves Glucose Tolerance in Zucker Diabetic Fatty (ZDF) Rats. Endocrinology. 2016;157(10):3844–55.

15. Chouchani ET, Kajimura S. Metabolic adaptation and maladaptation in adipose tissue. Nat Metab. 2019;1(2):189–200.

16. Hodson L, Gunn PJ. The regulation of hepatic fatty acid synthesis and partitioning: the effect of nutritional state. Nat Rev Endocrinol. 2019;15(12):689–700.

17. Grzybek M, Palladini A, Alexaki VI, Surma MA, Simons K, Chavakis T, et al. Comprehensive and quantitative analysis of white and brown adipose tissue by shotgun lipidomics. Mol Metab. 2019;22:12–20.

18. Vegiopoulos A, Rohm M, Herzig S. Adipose tissue: between the extremes. EMBO J. 2017;36(14):1999–2017.

19. Nam M, Choi MS, Jung S, Jung Y, Choi JY, Ryu DH, et al. Lipidomic Profiling of Liver Tissue from Obesity-Prone and Obesity-Resistant Mice Fed a High Fat Diet. Sci Rep. 2015;5:16984.

20. Eisinger K, Krautbauer S, Hebel T, Schmitz G, Aslanidis C, Liebisch G, et al. Lipidomic analysis of the liver from high-fat diet induced obese mice identifies changes in multiple lipid classes. Exp Mol Pathol. 2014;97(1):37–43.

21. Wu CL, Kimmerling KA, Little D, Guilak F. Serum and synovial fluid lipidomic profiles predict obesity-associated osteoarthritis, synovitis, and wound repair. Sci Rep. 2017;7:44315.

22. Eisinger K, Liebisch G, Schmitz G, Aslanidis C, Krautbauer S, Buechler C. Lipidomic analysis of serum from high fat diet induced obese mice. Int J Mol Sci. 2014;15(2):2991–3002.

23. Chatzigeorgiou A, Chung KJ, Garcia-Martin R, Alexaki VI, Klotzsche-von Ameln A, Phieler J, et al. Dual role of B7 costimulation in obesity-related nonalcoholic steatohepatitis and metabolic dysregulation. Hepatology. 2014;60(4):1196–210.

24. Chung KJ, Chatzigeorgiou A, Economopoulou M, Garcia-Martin R, Alexaki VI, Mitroulis I, et al. A self-sustained loop of inflammation-driven inhibition of beige adipogenesis in obesity. Nat Immunol. 2017;18(6):654–64.

25. Garcia-Martin R, Alexaki VI, Qin N, Rubin de Celis MF, Economopoulou M, Ziogas A, et al. ADIPOCYTE-SPECIFIC HIF2alpha DEFICIENCY EXACERBATES OBESITY-INDUCED BROWN ADIPOSE TISSUE DYSFUNCTION AND METABOLIC DYSREGULATION. Mol Cell Biol. 2015.

26. Sampaio JL, Gerl MJ, Klose C, Ejsing CS, Beug H, Simons K, et al. Membrane lipidome of an epithelial cell line. Proc Natl Acad Sci U S A. 2011;108(5):1903–7.

27. Ejsing CS, Sampaio JL, Surendranath V, Duchoslav E, Ekroos K, Klemm RW, et al. Global analysis of the yeast lipidome by quantitative shotgun mass spectrometry. Proc Natl Acad Sci U S A. 2009;106(7):2136–41.

28. Surma MA, Herzog R, Vasilj A, Klose C, Christinat N, Morin-Rivron D, et al. An automated shotgun lipidomics platform for high throughput, comprehensive, and quantitative analysis of blood plasma intact lipids. Eur J Lipid Sci Technol. 2015;117(10):1540–9.

29. Liebisch G, Binder M, Schifferer R, Langmann T, Schulz B, Schmitz G. High throughput quantification of cholesterol and cholesteryl ester by electrospray ionization tandem mass spectrometry (ESI-MS/MS). Biochim Biophys Acta. 2006;1761(1):121–8.

30. Herzog R, Schuhmann K, Schwudke D, Sampaio JL, Bornstein SR, Schroeder M, et al. LipidXplorer: a software for consensual cross-platform lipidomics. PLoS One. 2012;7(1):e29851.

31. Herzog R, Schwudke D, Schuhmann K, Sampaio JL, Bornstein SR, Schroeder M, et al. A novel informatics concept for high-throughput shotgun lipidomics based on the molecular fragmentation query language. Genome Biol. 2011;12(1):R8.

32. Pamplona R, Portero-Otin M, Riba D, Ruiz C, Prat J, Bellmunt MJ, et al. Mitochondrial membrane peroxidizability index is inversely related to maximum life span in mammals. J Lipid Res. 1998;39(10):1989–94.

33. R Core Team. R: A language and environment for statistical computing. 2018.

34. Wickham H. ggplot2: Elegant Graphics for Data Analysis.. Springer-Verlag New York. 2016;https://ggplot2.tidyverse.org

35. Peitzsch M, Dekkers T, Haase M, Sweep FC, Quack I, Antoch G, et al. An LC-MS/MS method for steroid profiling during adrenal venous sampling for investigation of primary aldosteronism. J Steroid Biochem Mol Biol. 2015;145:75–84.

36. Chatzigeorgiou A, Seijkens T, Zarzycka B, Engel D, Poggi M, van den Berg S, et al. Blocking CD40-TRAF6 signaling is a therapeutic target in obesity-associated insulin resistance. Proc Natl Acad Sci U S A. 2014;111(7):2686–91.

37. Yang Y, Lee M, Fairn GD. Phospholipid subcellular localization and dynamics. J Biol Chem. 2018;293(17):6230–40.

38. van der Veen JN, Kennelly JP, Wan S, Vance JE, Vance DE, Jacobs RL. The critical role of phosphatidylcholine and phosphatidylethanolamine metabolism in health and disease. Biochim Biophys Acta Biomembr. 2017;1859(9 Pt B):1558–72.

39. Sunshine H, Iruela-Arispe ML. Membrane lipids and cell signaling. Curr Opin Lipidol. 2017;28(5):408–13.

40. van Meer G, Voelker DR, Feigenson GW. Membrane lipids: where they are and how they behave. Nat Rev Mol Cell Biol. 2008;9(2):112–24.

41. Pergande MR, Serna-Perez F, Mohsin SB, Hanek J, Cologna SM. Lipidomic Analysis Reveals Altered Fatty Acid Metabolism in the Liver of the Symptomatic Niemann-Pick, Type C1 Mouse Model. Proteomics. 2019;19(18):e1800285.

42. Sonnweber T, Pizzini A, Nairz M, Weiss G, Tancevski I. Arachidonic Acid Metabolites in Cardiovascular and Metabolic Diseases. Int J Mol Sci. 2018;19(11).

43. Hanna VS, Hafez EAA. Synopsis of arachidonic acid metabolism: A review. J Adv Res. 2018;11:23–32.

44. Nakamura MT, Nara TY. Structure, function, and dietary regulation of delta6, delta5, and delta9 desaturases. Annu Rev Nutr. 2004;24:345–76.

45. Brash AR. Arachidonic acid as a bioactive molecule. J Clin Invest. 2001;107(11):1339–45.

46. Pasquali R, Cantobelli S, Casimirri F, Capelli M, Bortoluzzi L, Flamia R, et al. The hypothalamic-pituitary-adrenal axis in obese women with different patterns of body fat distribution. J Clin Endocrinol Metab. 1993;77(2):341–6.

47. Wallerius S, Rosmond R, Ljung T, Holm G, Bjorntorp P. Rise in morning saliva cortisol is associated with abdominal obesity in men: a preliminary report. J Endocrinol Invest. 2003;26(7):616–9.

48. Qi D, Rodrigues B. Glucocorticoids produce whole body insulin resistance with changes in cardiac metabolism. Am J Physiol Endocrinol Metab. 2007;292(3):E654–67.

49. Montgomery MK, Brown SHJ, Mitchell TW, Coster ACF, Cooney GJ, Turner N. Association of muscle lipidomic profile with high-fat diet-induced insulin resistance across five mouse strains. Sci Rep. 2017;7(1):13914.

50. Cheng B, Kowal J. Analysis of adrenal cholesteryl esters by reversed phase high performance liquid chromatography. J Lipid Res. 1994;35(6):1115–21.

51. Vance JE. Historical perspective: phosphatidylserine and phosphatidylethanolamine from the 1800s to the present. J Lipid Res. 2018;59(6):923–44.

52. Testerink N, van der Sanden MH, Houweling M, Helms JB, Vaandrager AB. Depletion of phosphatidylcholine affects endoplasmic reticulum morphology and protein traffic at the Golgi complex. J Lipid Res. 2009;50(11):2182–92.

53. Patel D, Witt SN. Ethanolamine and Phosphatidylethanolamine: Partners in Health and Disease. Oxid Med Cell Longev. 2017;2017:4829180.

54. Furse S, de Kroon AI. Phosphatidylcholine’s functions beyond that of a membrane brick. Mol Membr Biol. 2015;32(4):117–9.

55. Cadas H, di Tomaso E, Piomelli D. Occurrence and biosynthesis of endogenous cannabinoid precursor, N-arachidonoyl phosphatidylethanolamine, in rat brain. J Neurosci. 1997;17(4):1226–42.

56. Fan W, Evans RM. Turning up the heat on membrane fluidity. Cell. 2015;161(5):962–3.

57. Hac-Wydro K, Wydro P. The influence of fatty acids on model cholesterol/phospholipid membranes. Chem Phys Lipids. 2007;150(1):66–81.

58. Holthuis JC, Menon AK. Lipid landscapes and pipelines in membrane homeostasis. Nature. 2014;510(7503):48–57.

59. Kraemer FB. Adrenal cholesterol utilization. Mol Cell Endocrinol. 2007;265–266:42–5.

60. Gallay J, Vincent M, de Paillerets C, Rogard M, Alfsen A. Relationship between the activity of the 3 beta-hydroxysteroid dehydrogenase from bovine adrenal cortex microsomes and membrane structure. Influence of proteins and steroid substrates on lipid “microviscosity”. J Biol Chem. 1981;256(3):1235–41.

61. Narasimhulu S. Thermotropic transitions in fluidity of bovine adrenocortical microsomal membrane and substrate-cytochrome P-450 binding reaction. Biochim Biophys Acta. 1977;487(2):378–87.

62. Stoffel W, Holz B, Jenke B, Binczek E, Gunter RH, Kiss C, et al. Delta6-desaturase (FADS2) deficiency unveils the role of omega3-and omega6-polyunsaturated fatty acids. EMBO J. 2008;27(17):2281–92.

63. Stroud CK, Nara TY, Roqueta-Rivera M, Radlowski EC, Lawrence P, Zhang Y, et al. Disruption of FADS2 gene in mice impairs male reproduction and causes dermal and intestinal ulceration. J Lipid Res. 2009;50(9):1870–80.

64. Schaeffer L, Gohlke H, Muller M, Heid IM, Palmer LJ, Kompauer I, et al. Common genetic variants of the FADS1 FADS2 gene cluster and their reconstructed haplotypes are associated with the fatty acid composition in phospholipids. Hum Mol Genet. 2006;15(11):1745–56.

65. Stoffel W, Hammels I, Jenke B, Binczek E, Schmidt-Soltau I, Brodesser S, et al. Obesity resistance and deregulation of lipogenesis in Delta6-fatty acid desaturase (FADS2) deficiency. EMBO Rep. 2014;15(1):110–20.

66. Roqueta-Rivera M, Stroud CK, Haschek WM, Akare SJ, Segre M, Brush RS, et al. Docosahexaenoic acid supplementation fully restores fertility and spermatogenesis in male delta-6 desaturase-null mice. J Lipid Res. 2010;51(2):360–7.

67. Zolfaghari R, Cifelli CJ, Banta MD, Ross AC. Fatty acid delta(5)-desaturase mRNA is regulated by dietary vitamin A and exogenous retinoic acid in liver of adult rats. Arch Biochem Biophys. 2001;391(1):8–15.

68. Vessby B. Dietary fat, fatty acid composition in plasma and the metabolic syndrome. Curr Opin Lipidol. 2003;14(1):15–9.

69. Warensjo E, Ohrvall M, Vessby B. Fatty acid composition and estimated desaturase activities are associated with obesity and lifestyle variables in men and women. Nutr Metab Cardiovasc Dis. 2006;16(2):128–36.

70. Kroger J, Schulze MB. Recent insights into the relation of Delta5 desaturase and Delta6 desaturase activity to the development of type 2 diabetes. Curr Opin Lipidol. 2012;23(1):4–10.

71. Kroger J, Zietemann V, Enzenbach C, Weikert C, Jansen EH, Doring F, et al. Erythrocyte membrane phospholipid fatty acids, desaturase activity, and dietary fatty acids in relation to risk of type 2 diabetes in the European Prospective Investigation into Cancer and Nutrition (EPIC)-Potsdam Study. Am J Clin Nutr. 2011;93(1):127–42.

72. Park H, Hasegawa G, Shima T, Fukui M, Nakamura N, Yamaguchi K, et al. The fatty acid composition of plasma cholesteryl esters and estimated desaturase activities in patients with nonalcoholic fatty liver disease and the effect of long-term ezetimibe therapy on these levels. Clin Chim Acta. 2010;411(21-22):1735–40.

73. Lopez-Vicario C, Gonzalez-Periz A, Rius B, Moran-Salvador E, Garcia-Alonso V, Lozano JJ, et al. Molecular interplay between Delta5/Delta6 desaturases and long-chain fatty acids in the pathogenesis of non-alcoholic steatohepatitis. Gut. 2014;63(2):344–55.

74. Kang JX, Wang J, Wu L, Kang ZB. Transgenic mice: fat-1 mice convert n-6 to n-3 fatty acids. Nature. 2004;427(6974):504.

75. Matias I, Petrosino S, Racioppi A, Capasso R, Izzo AA, Di Marzo V. Dysregulation of peripheral endocannabinoid levels in hyperglycemia and obesity: Effect of high fat diets. Mol Cell Endocrinol. 2008;286(1-2 Suppl 1):S66–78.

76. Kurahashi Y, Ueda N, Suzuki H, Suzuki M, Yamamoto S. Reversible hydrolysis and synthesis of anandamide demonstrated by recombinant rat fatty-acid amide hydrolase. Biochem Biophys Res Commun. 1997;237(3):512–5.

77. Willoughby KA, Moore SF, Martin BR, Ellis EF. The biodisposition and metabolism of anandamide in mice. J Pharmacol Exp Ther. 1997;282(1):243–7.

78. Byrd JB, Turcu AF, Auchus RJ. Primary Aldosteronism: Practical Approach to Diagnosis and Management. Circulation. 2018;138(8):823–35.

79. Loriaux DL. Diagnosis and Differential Diagnosis of Cushing’s Syndrome. N Engl J Med. 2017;377(2):e3.

