## Supplementary data for "Obesity-associated lipidomic remodeling of the adrenal gland indicates an important role of the FADS2-arachidonic acid axis in adrenocortical hormone production"

**Supplementary Table 1. List of primers**

| | |
|---|---|
| *Cyp11a1*_Fwd | GCTGAGTACTGGAAAGGGAGC |
| *Cyp11a1*_Rev | TGCCCAGCTTCTCCCTGTAA |
| *Cyp11b1*_Fwd | ATAGAGAACTCCGTGGCCTG |
| *Cyp11b1*_Rev | TGCAGTCGGTTGAAGTACCA |
| *Cyp11b2*_Fwd | AGAATGGCGTCTCAACCGAC |
| *Cyp11b2*_Rev | AGTCCCTTGCTACCATGTCC |
| *Cyp21a1*_Fwd | GCTGTGGCTTTCCTGCTTCAC |
| *Cyp21a1*_Rev | GGCCCAGCTTGAGGTCTAACT |
| *Fads2*_Fwd | TCCTCTCGTACTTCGGCACT |
| *Fads2*_Rev | GGTTCCACCAGTTGGCTGAG |
| *Hsd3b1*_Fwd | GCCACCACCATCTCAGACTTT |
| *Hsd3b1*_Rev | GGTAACCCTTAGGAGGGCTGTG |
| *Hsd3b2*_Fwd | AGGGAGCTCTCAATTGTGCC |
| *Hsd3b2*_Rev | GCTTAGAAAGGCTGGTTCTGG |
| *Star*_Fwd | CTGTCCACCACATTGACCTG |
| *Star*_Rev | CAGCTATGCAGTGGGAGACA |
| *Tbp*_Fwd | AGAACAATCCAGACTAGCAGCA |
| *Tbp*_Rev | GGGAACTTCACATCACAGCTC |

**Supplementary Table 2. DBI and LI in storage and membrane lipids in the adrenal gland from ND- and HFD-fed mice.**

| | **STORAGE LIPIDS** | | | **MEMBRANE LIPIDS** | | |
|---|---|---|---|---|---|---|
| | **ND** | **HFD** | **P-value** | **ND** | **HFD** | **P-value** |
| **DBI** | 2.809 +/- 0.095 | 3.072 +/- 0.024 | *** | 3.322 +/- 0.049 | 3.462 +/- 0.042 | *** |
| **LI** | 45.291 +/- 4.203 | 47.537 +/- 1.223 | ns | 36.828 +/- 0.081 | 37.110 +/- 0.064 | *** |

**Supplementary Figure 1. Lipid class composition of the adrenal gland lipidome in lean and obese mice**

Mean mol % ± standard deviation of all lipid classes detected in adrenal glands from mice fed for 20 weeks a ND or a HFD. n=7-8. The upper panel shows the predominant classes, the lower panel the classes of low abundance. Mol % is relative to the whole lipidome.

**Supplementary Figure 2. Lipid length and unsaturation profile of storage and non-storage lipids in the adrenal glands of lean and obese mice**

**A,C.** Storage (CE, TAG) (**A**) and non-storage lipids (phospholipids, lysophospholipids and sphingolipids; cholesterol was excluded from the calculation) (**C**) in adrenal glands of mice fed for 20 weeks a ND or HFD were regrouped according to the number of carbon atoms of the acyl chains; only the length groups > 1 mol % are shown. **B,D**. Storage (**B**) and non-storage lipids (**D**) were regrouped according to the number of double bonds of their acyl chains. The mean mol % ± standard deviation are shown, n=7-8; * p-value<0.05; ** p-value<0.01; *** p-value<0.001.

**Supplementary Figure 3. Gene expression of steroidogenic enzymes in adrenal cell cultures treated with FADS2 inhibitor**

Relative mRNA expression of *Star, Cyp11a1, 3bHsd2, Cyp21a1, Cyp11b1* and *Cyp11b2* in primary adrenal gland cell cultures treated for 18 h with FADS2 inhibitor (FADS2i) or DMSO (Ctrl). *Tbp* expression was used as an internal control. Relative gene expression of DMSO-treated cells was set to 1. Data are shown as mean ± SEM; n=12; ** p-value<0.01, *** p-value<0.001; **** p-value<0.0001.

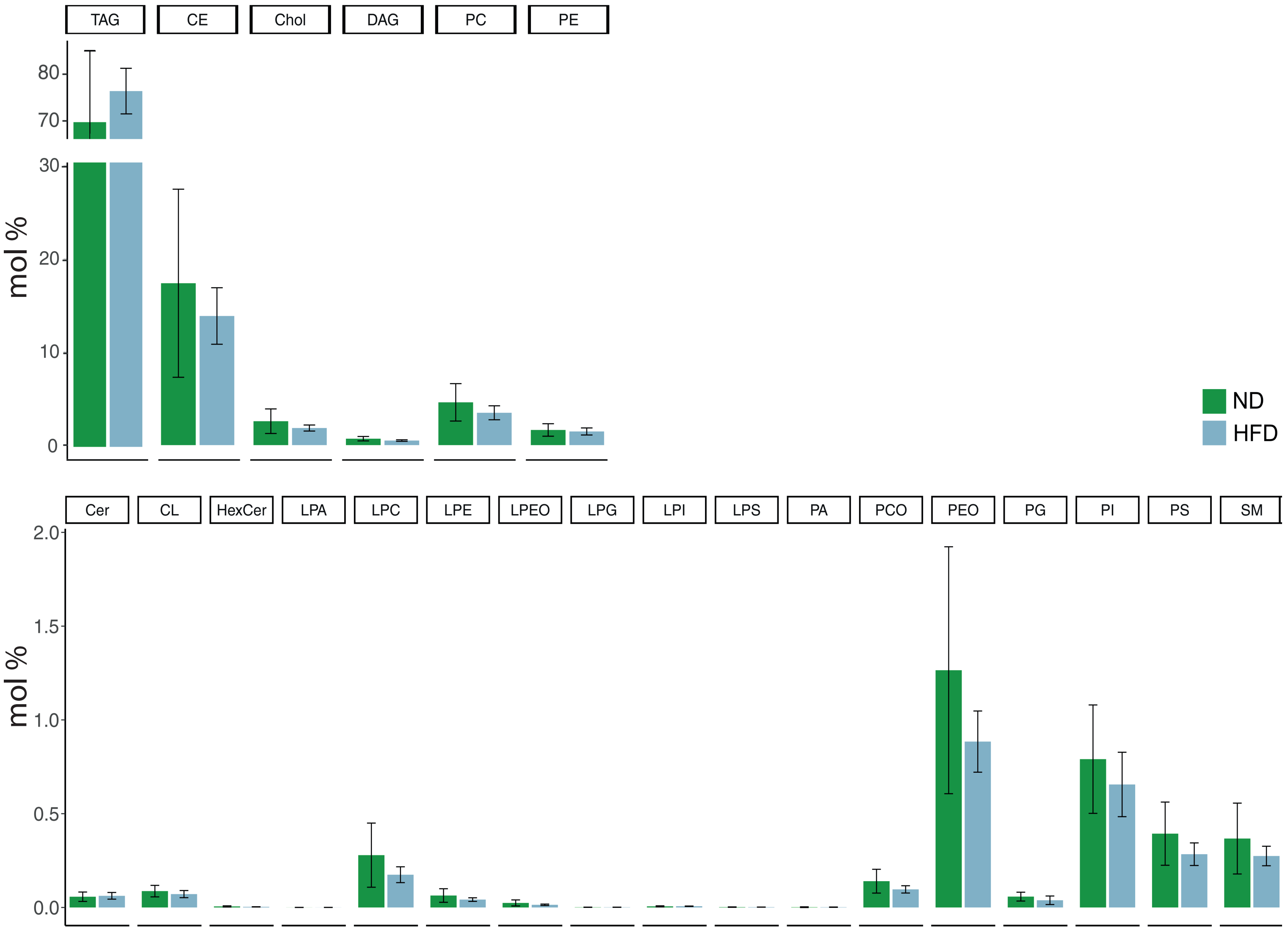


Supplementary Figure 1

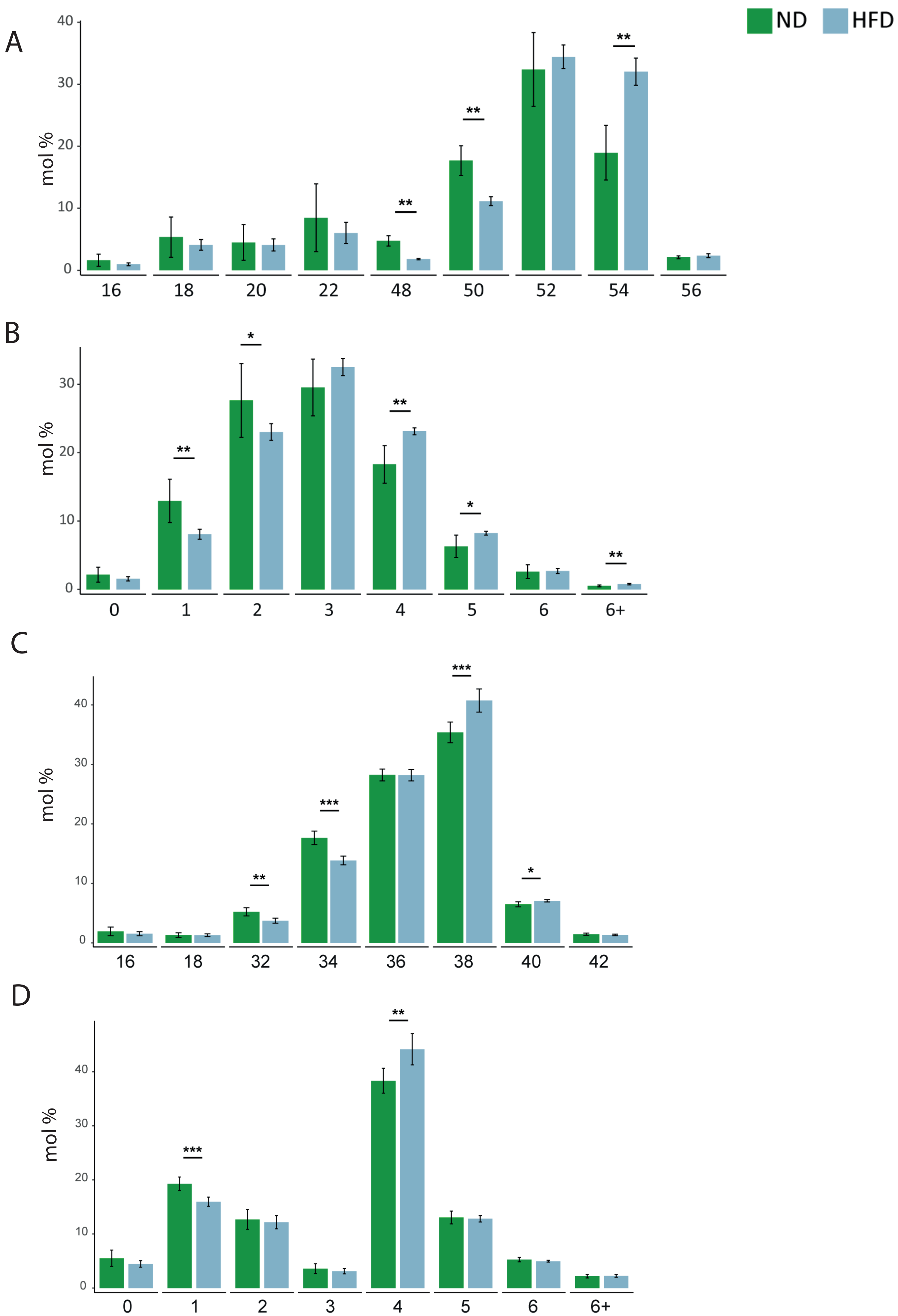


Supplementary Figure 2

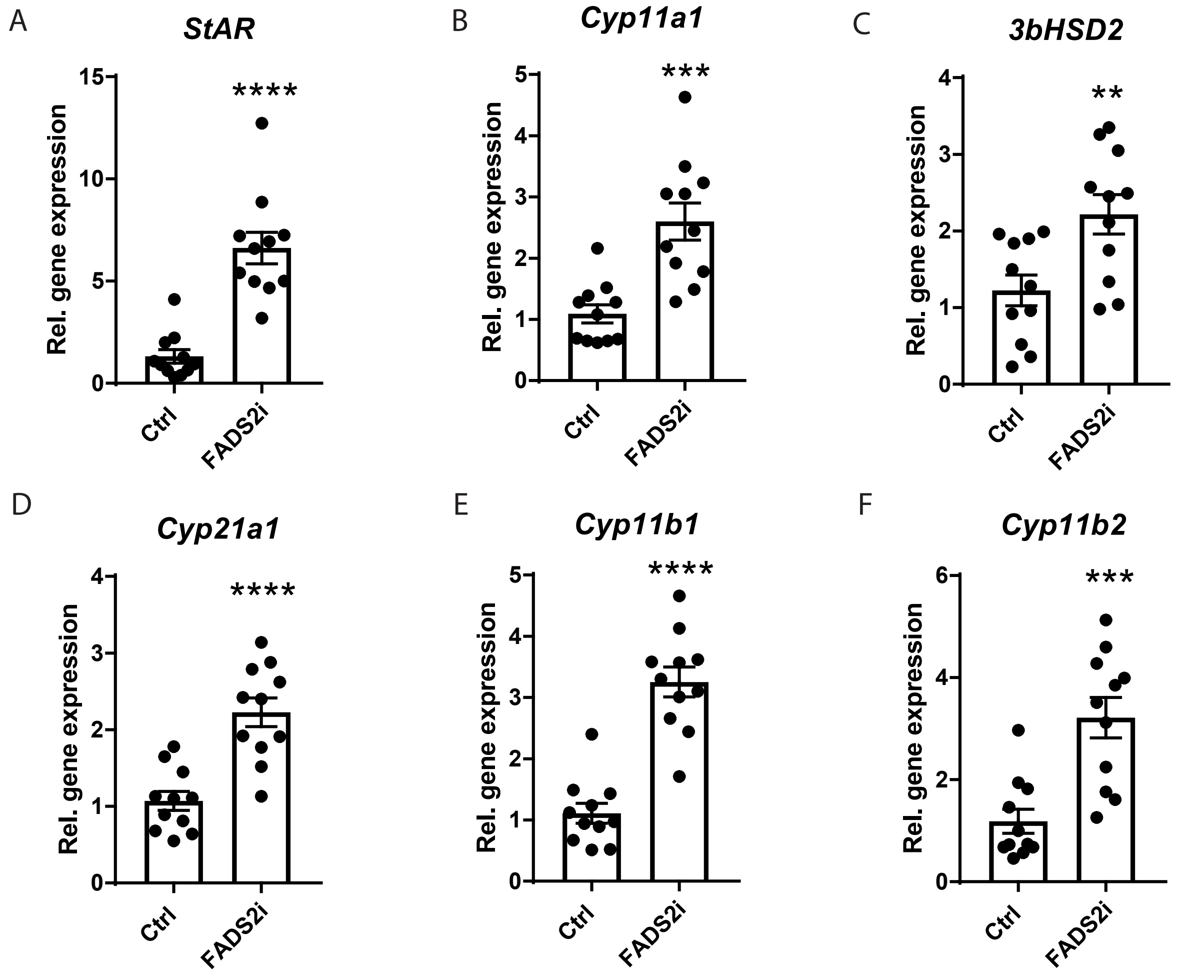


Supplementary Figure 3
